# *De novo* Genome Assembly, Functional Annotation and SSR Mining of *Citrus reticulata* “Kinnow” from Pakistan

**DOI:** 10.1101/2023.03.27.534305

**Authors:** Sadia Jabeen, Rashid Saif, Gaetano Distefano, Rukhama Haq, Waseem Haider, Akber Hayat, Shagufta Naz

## Abstract

*Citrus reticulata* (Blanco) fruit is native to South East Asia which owns many of the nutritional, medicinal and economic advantages, locally known as “Kinnow” and one of the priced mandarin varieties (Dancy, Fuetrell’s Early and Honey) of *Citrus* genera renowned for its exclusive taste, vitamin richness, thin peel, long shelf-life and seedless characteristics in Pakistan. However, genetic improvement and breeding strategies of this valued variety are lacking due to the in-housed insufficient genomic and technical resources. Therefore, the current research was initiated to provide the base-line *de-novo* genome assembly of *C. reticulata* (seedless kinnow) at a depth of 151x with Illumina paired-end short-read sequencing technology using HiSeq 2500. Whole-genome sequencing resulted in 139,436,350 raw reads (∼20.09 GB) of data, however, after removing the low-quality reads (1.08%), duplicated sequences (10.5%) and Illumina adaptors, 137,901,462 clean reads were obtained with (∼18.87 GB) of clean data which was further used for downstream variant calling analysis. In total, 348,861 scaffolds were generated with N50 value of 4827 which constitute 263,018,9 contigs ranging from 71-36,213 with total of 179,984,763 nucleotides. The GC content of the final draft assembly at 71-mer was 34.1%. Moreover, annotation was performed with “Hayai-Annotation Plants” tool which marked the whole-genome mapping with three main functional databases of interpro, Pfam and gene ontology. Additionally, in-silico identification of 111,032 Simple Sequence Repeats (SSR) was also accomplished with the help of GMATA tool, which may be used for further screening and genetic improvement of the citrus varieties by means of this current assembly as a resource of local reference genome.

## Introduction

The genus Citrus comprises numerous economically important species within the Rutaceae family (Appelhans *et al*. 2021), which are grown in tropical and temperate regions worldwide (∼159 countries) with a total production of 158 million tons (FAO statistics. 2021). Citrus fruits thrive in regions with tropical, subtropical, or mixed climates. Although the birthplace of citrus is uncertain, Southeast Asia is believed to be its origin. The taxonomy and phylogeny of citrus are challenging due to various factors such as bud mutations rate, the ability of citrus and related genera to interbreed, widespread distribution, and a long history of cultivation (Shahnazari *et al*., 2022).

The principal species of the genus Citrus are *C. reticulata* (mandarin), *C. maxima* (Pummelo), and *C. medica* (Citron) (Wu *et al*., 2018), while other species such as *C. aurantium* (sour orange), *C. sinensis* (sweet orange), *C. paradisi* (grapefruit), and *C. limon* (lemon) are derived from successive hybridizations (Mario *et al*., 2021). The citrus genome is typically diploid, consisting of nine chromosomes (Krug. 1943), but other euploid genotypes such as triploid *F. hindsii* (kumquat), *C. latifolia* (Tahiti lime), and tetraploid *P. trifoliata* are also studied (Wu *et al*., 2018).

As of 2020, mandarin production ranked second after sweet orange, accounting for 24% of worldwide citrus production (FAO statistics. 2020), and ranked first in Pakistan (Raza et al. 2021). Nowadays, mandarin fruits are gaining attention due to their appealing attributes such as a nutritional profile, ease of peeling, and unique appetizing flavor. The well-known commercial varieties of mandarin include Satsuma mandarin, Clementine mandarin, and local landraces, including Dancy, Feutrells Early, and Kinnow in Pakistan (Usman and Fatima. 2018).

The Kinnow mandarin is hybrid of King and Willow leaf (Kaur *et al*., 2022). It is native to California as in 1915 H. B. Frost developed Kinnow first time at the University of California Research Center, Riverside. Almost after 20 years later, in 1935, it was released to commercial market including Pakistan (Mahawar *et al*., 2020). It is grown in all four provinces of Pakistan, with Punjab responsible for over 95% of all crops due to ideal environmental conditions for growth, including enough water, temperature and soil. Pakistan produces 6.5% of the world’s mandarins, 60% of which are Kinnow and 5% are other varieties. As the sixth-ranked mandarin grower globally, Pakistan’s estimated 370 thousand tonnes of production have a 222-million-dollar export value (Nawaz *et al*., 2018).

In Pakistan, two varieties of Kinnow are cultivated: seeded and seedless. Both types share some similarities and differences. Morphologically, they are similar in tree shape, branch angle, vegetative life cycle, leaf apex, lamina, leaf blade, length of petiole, scion trunk surface, fruit segment shape, and albedo color. However, seedless Kinnow is lighter in weight than the seeded variety, owing to the absence of seeds. Additionally, PCA analysis also indicated that weight of fruit, peel, fruit diameter, and non-reducing sugars were the major contributing factors for variations between the two varieties. Furthermore, genetic differences between the two varieties are also confirmed by RAPD and SSR markers (Altaf *et al*., 2014).

Seeded Kinnow is highly nutritious and in high demand, but its high number of seeds per fruit makes it a low-value export fruit (Khalil et al. 2011). In contrast, seedless Kinnow has several advantages over seeded Kinnow, including gustatory benefits, better taste, greater sweetness due to a significant number of soluble solids, and benefits for the citrus fruit processing industry as seeds do not need to be removed during processing. The cavities that contain the seeds in seeded Kinnow are replaced with edible tissues in seedless Kinnow, which is more appealing to consumers. Furthermore, seedless Kinnow has a longer shelf life compared to seeded Kinnow, which has a shorter shelf life due to the hormones produced by the seeds that trigger senescence. Additionally, seedless Kinnow contains higher levels of ascorbic acid, citric acid, malic acid, hesperidin, flavonoids, and β-carotene than seeded Kinnow (Premachandran *et al*., 2019 ; Jagga *et al*., 2022).

The field of citrus genomics has seen substantial growth in recent years, thanks to advancements in sequencing technology. With the significant reduction in the cost of high-throughput sequencing, there has been a sharp increase in the amount of DNA sequence data available for analysis. Among the citrus species whose genomes have been published are pummelo, citron, clementine, and mandarin (Wang *et al*., 2017). This wealth of genomic data has greatly benefitted citrus researchers and breeders, as it allows for the identification of desirable traits and the development of improved cultivars. Moreover, the genomic data has contributed to a better understanding of the evolutionary history of the genus Citrus and has provided deeper insight into the genetic basis of important agronomic traits. These advancements in citrus genomics offer exciting prospects for the future of citrus breeding and improvement.

The identification of various citrus species has been achieved through the use of molecular techniques such as Simple Sequence Repeat (SSR) markers. Research indicates that the majority of mandarin cultivars exhibit limited genetic diversity (Kaur *et al*., 2022), likely due to somatic mutations that cannot be detected by some molecular markers. SSR markers are a reliable tool for fingerprinting and analyzing genetic variation in mandarin and other citrus species. However, they are limited by homoplasy, a phenomenon in which the same SSR marker can rate DNA fragments based on size (Shahnazari *et al*., 2022).

Given the medicinal and economic value of seedless Kinnow, its whole genome was sequenced and annotated to identify potential genes that could be further exploited to enhance economically significant traits. Annotation was performed with the “Hayai-Annotation Plants” tool, which marked the whole genome mapping with three main functional databases: interpro, Pfam, and gene ontology. Additionally, in-silico identification of 111,032 SSRs was achieved using the GMATA tool, which could be further employed for screening and genetic improvement of local citrus varieties

## Materials and Methods

### Sample collection, DNA extraction and quantification

The fresh and healthy leaves were collected from the single true-to-type plant of kinnow cultivated in Citrus Research Institute Sargodha and the total genomic DNA was extracted using the Murray and Thompson (1980) protocol with some modifications (Naz *et al*., 2014). The extracted DNA was stored at -20°C (Fig.1 A). Nanodrop™ (Thermo Scientific, Germany) was used to check the purity and concentration of the DNA. Before sequencing, the DNA samples were checked for contamination and DNA degradation by agarose gel electrophoresis and the DNA concentration was quantified by Qubit 2.0 (Liu *et al*., 2022).

**Figure 1:**
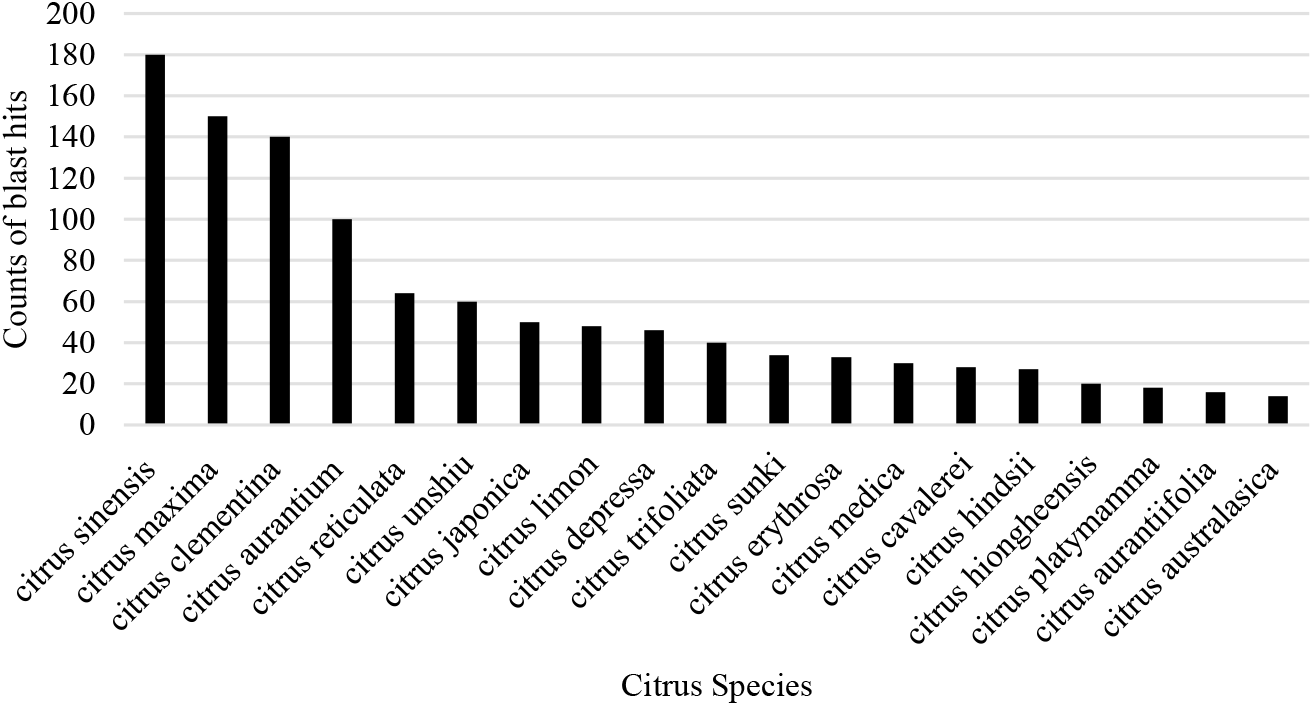
Top hits in Nr databases. y-axis represents the number of hits against the Citrus species on x-axis

### Next-generation Illumina sequencing

Sequencing libraries were prepared using the NEBNext® DNA Library preparation kit. Random shearing of DNA produced countless fragments consist of 350 bp, which are ligated with NEBNext® adapter, and amplified with P5 and indexed P7 oligos for PCR. AMPure XP system was used for purification of amplified libraries. On the Illumina HiSeq 2500 the paired-end (2×150) libraries were pooled and sequenced.

### In-silico quality checks of the raw reads

Fastq files were processed before proceeding with data for downstream analysis with FASTQC (v.0.119) (Andrews, 2010). All the quality checks were performed including per base sequence quality, per tile sequence quality, per sequence quality score, per base sequence content, per sequence GC content, per base N content, and sequence length distribution, sequence duplication level, and adaptor content.

### Whole-genome de-novo assembly with platanus & curation

As citrus is a highly heterozygous species (Kundu and Dubey. 2022), the genome assembly was performed with the Platanus software (v.1.2.4) as previous works described its reliability in the assembly of highly heterozygous genomes (Patel *et al*., 2018 ; Habibi *et al*., 2022). Platanus worked based on three basic steps a) assembly, b) scaffolding and c) gap closing. For assembly the clean trimmed sequences were assembled to generate contigs, followed by scaffolding and gap closing ultimately generate the consensus sequence. QUAST was used for assembly statistics (Skóra *et al*., 2022). This tool comprises different assessment metrics, such as assemblies, contig sizes, structural variations, N50 and genomic functional elements. The annotation was performed using BLASTx against the Nr (NCBI non-redundant) database with a cut-off E-value set to 10^−5^. Results were further verified with another comprehensive and ultrafast tool: Hayai-Annotation Plants, for functional annotation with default parameters (Ghelfi *et al*., 2019). This tool is specifically designed for the *in-silico* annotation of plants consist of three main and steps (a) input of the Fasta files, (b) Sequence alignment (c) Functional annotation. The evolutionary history was inferred by using the Maximum Likelihood method and Tamura-Nei model in MEGA11 (Tamura *et al*., 2021)

### Identification of microsatellite/SSR motif

For the efficient mining of microsatellite motifs GMATA (v 2.0) was used with default parameters (Wang and Wang. 2016) The usual parameters like primer size: 18-23 bp; annealing temperature: 57-62°C; GC content: 30-70%; end product size: 100-400 bp were utilized to build primers against filtered SSR motifs

## Results

### Detailed statistics for the quality checks of sequencing data

The accuracy of Illumina sequencing platform can be assessed by the Q score (Phred quality score), this metric is used to measure the probability of base calling to assure the accuracy and authenticity of the called base position. Generally, Q30 score means no ambiguity and zero error in base calling, that’s why it is considered as a criterion for quality in next-generation sequencing. All the phred quality scores with GC % are given in Supplementary Table S1.

### Basic statistics of de-novo genome assembly

The total data generate on flow cell as R1 was 3.998.020 kb and R2 as 4,386,275 kb. On second flow cell the total sequences were 1,122,009 kb and R2 was 1,195,659 kb, and for the third flow cell the R1 was observed as 739,159 kb and R2 as 833,588 kb. Genome sequencing resulted in 139,436,350 raw reads (∼20.09 GB) of data, after removing the low-quality reads (1.08%), duplicated sequences (10.5%) and Illumina adaptors, 137,901,462 clean reads were obtained (∼18.87 GB) that were further used for the downstream analysis. In total, 348,861 scaffolds were generated with N50 value of 4,827 which constitute 2,630,18,9 contigs with largest contig size 36,213 with total 179,984,763 nucleotides. Table 1 highlighted details statistics of assembly. The GC content of the final draft assembly at 71-mer was 34.1%. Based on the results, we can say that the *de novo* assembly is complicated due to its complexity.

**Table 1:** General statistics for *Citrus reticulata* genome assembly using Platanus.

| Assembly statistics without reference | Contig (71-mers) | Scaffold (71-mers) | Gap closed (71-mers) |
| --- | --- | --- | --- |
| Contigs | 137054 | 93403 | 93532 |
| # contigs ($\geq 0$ bp) | 2630189 | 348861 | 348861 |
| # contigs ($\geq 1000$ bp) | 52379 | 57276 | 57383 |
| # contigs ($\geq 5,000$ bp) | 3896 | 11267 | 11273 |
| # contigs ($\geq 10,000$ bp) | 693 | 4010 | 4007 |
| # contigs ($\geq 25,000$ bp) | 22 | 478 | 479 |
| # contigs ($\geq 50,000$ bp) | 0 | 55 | 55 |
| Largest contig/scaffold | 36213 | 114892 | 114879 |
| Total length | 179984763 | 243652840 | 243869956 |
| Total length ($\geq 0$ bp) | 514402379 | 289058864 | 289231328 |
| Total length ($\geq 1000$ bp) | 122227900 | 218060465 | 218263338 |
| Total length ($\geq 5,000$ bp) | 30894127 | 119069997 | 119117432 |
| Total length ($\geq 10,000$ bp) | 9722230 | 68967661 | 68944761 |
| Total length ($\geq 25,000$ bp) | 634487 | 17254302 | 17277800 |
| Total length ($\geq 50,000$ bp) | 0 | 3637106 | 3637277 |
| N50 | 1614 | 4827 | 4819 |
| N90 | 608 | 973 | 973 |
| L50 | 26594 | 11829 | 11848 |
| L90 | 104305 | 58521 | 58620 |
| GC (%) | 34.1 | 34 | 34 |

### Phylogenetics of assembled genome

To understand the similarity with the other citrus verities and the present genome sequence alignment using Blast2Sequence was performed. The blast hits distribution in the Nr database showed most hits with *Citrus sinensis*. The second top blast hit species was *Citrus maxima* (Fig. 1).

The tree with the highest log likelihood (−1905.05) is shown (Fig.2). The percentage of trees in which the associated taxa clustered together is highlighted next to the branches. Initial tree(s) for the heuristic search were obtained automatically by applying Neighbor-Join and BioNJ algorithms to a matrix of pairwise distances estimated using the Tamura-Nei model, and then selecting the topology with superior log likelihood value. This analysis involved 28 nucleotide sequences. There were a total of 1128 positions in the final dataset. Current analysis suggesting that the genomic contents of seedless Kinnow might be similar to those of the seeded Kinnow.

**Figure 2:**
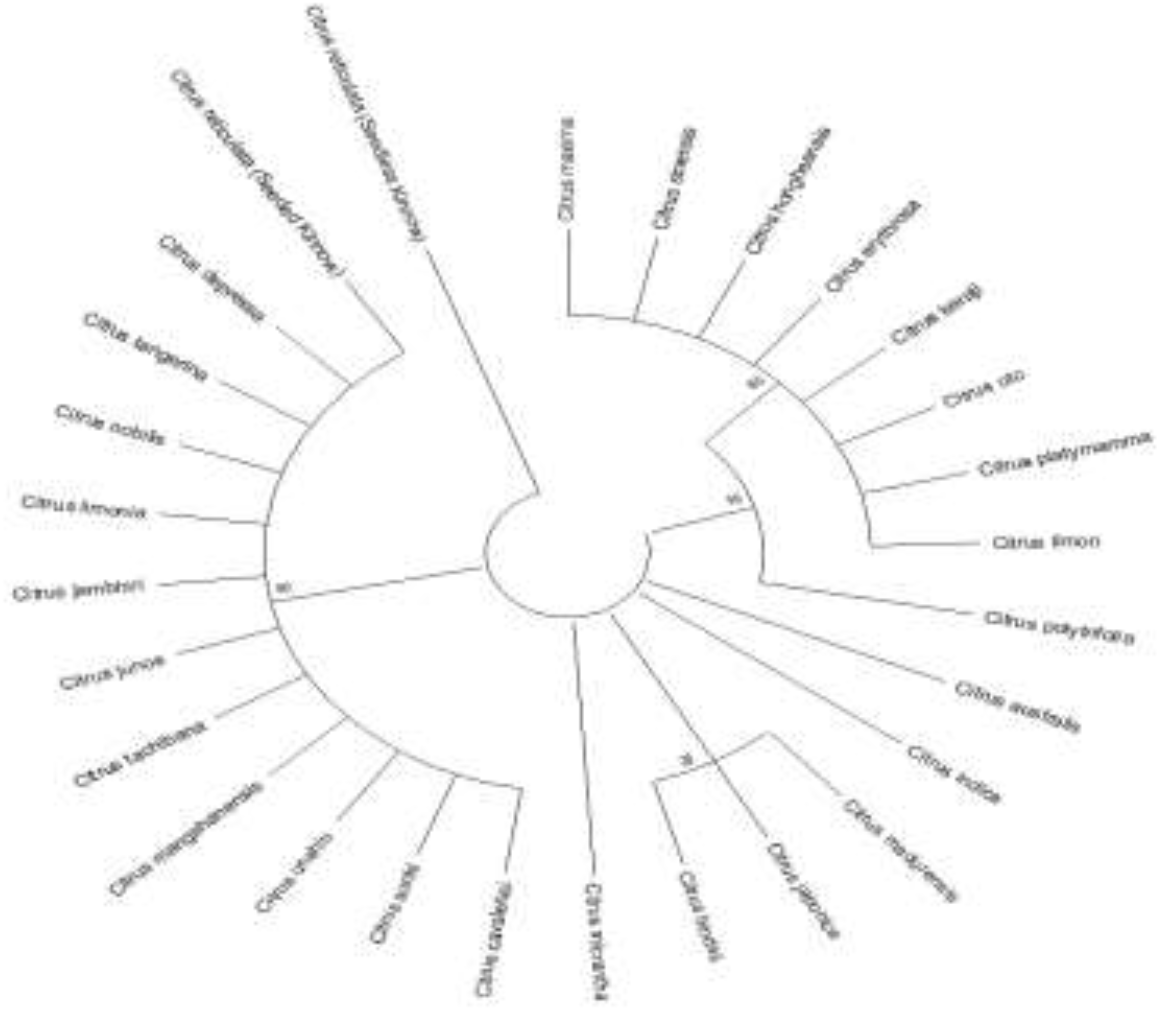
Dendrogram represents the evolutionary relationship between seedless Kinnow and other citrus species

### Functional annotation of assembled genome

All the assembled contigs from our studied genome of *Kinnow* were annotated by Hayai-Annotation-Plants platform. It generates gene annotation with accuracy for the family members of Embryophyta (plant species). Here it produced annotation for three main categories: interpro, Pfam, gene ontology (biological processes GO:BP, cellular components GO:CC, molecular functions GO:MF). It compared our reported contigs with databases. The analysis resulted in 1029 contigs showing similarities with interpro terms (Fig 3a), 527 with Pfam (Fig 3b), 302 contigs annotated as GO:MF (Fig 3c), 123 with GO:BP (Fig 3d), and 91 with GO:CC (Fig 3e). Surprisingly, interpro and Pfam both top most annotated term was RVT-2 (conserved domain of reverse transcriptase) which is involved in the different biochemical activities at cellular level and in genetic diversity (Supplementary Fig. 1 and 2). DNA integration term (66%) in GO:BP were observed as top most on the basis of annotations from different databases (Supplementary Fig. 3). Nucleic acid binding genes (93%) in GO:MF (Supplementary Fig. 4)., while integral component of cell membrane (41%) in GO:CC (Supplementary Fig. 5). Additionally, the detailed results of annotated assembled whole genome are given in Supplementary Tables (S2 to S7).

**Figure 3:**
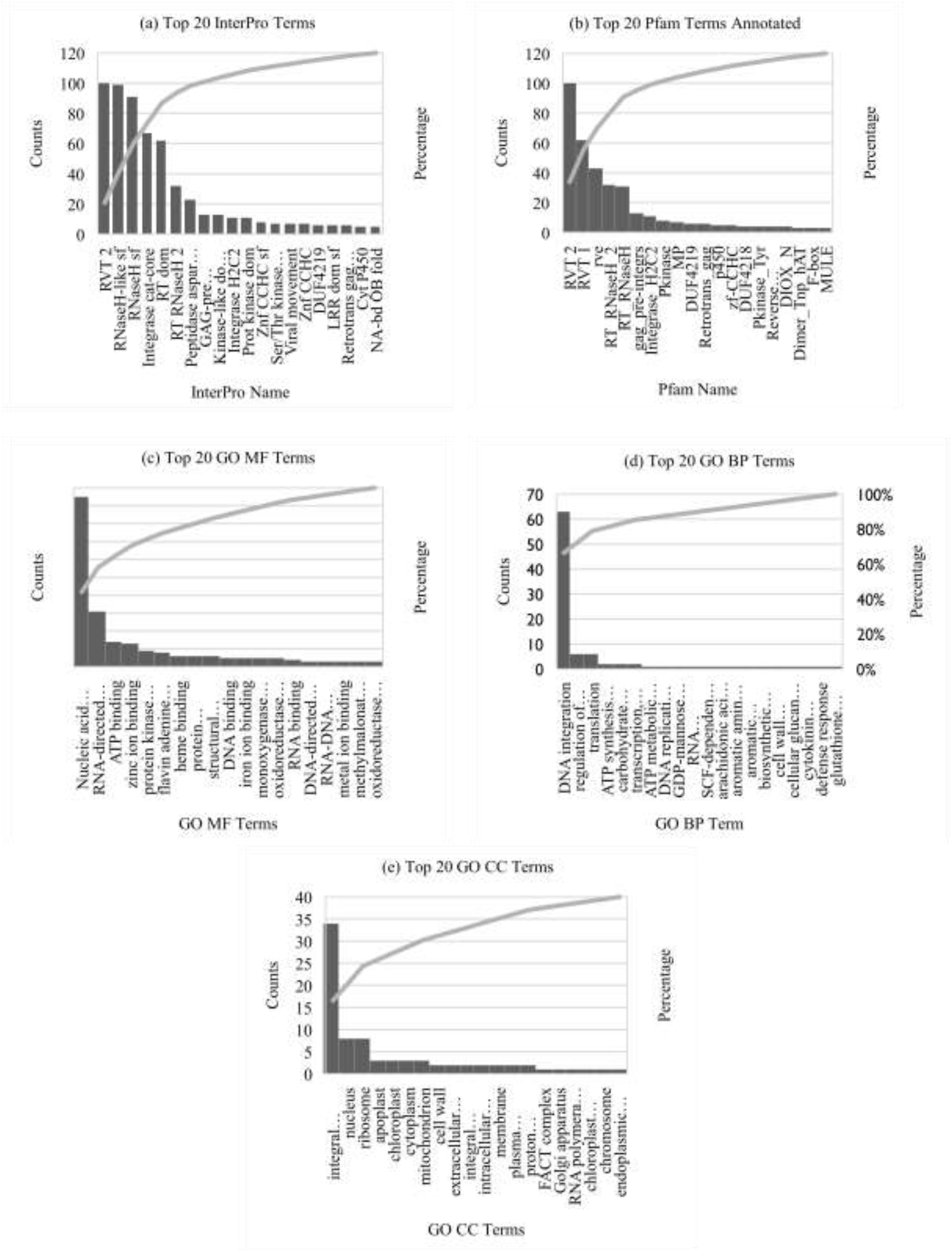
Hayai-annotation results (a) Top 20 annotated terms from Interpro (b) Top 20 annotated terms from Pfam (c) Top 20 annotated terms from gene ontology molecular functions (d) Top 20 annotated terms from gene ontology biological processes (e) Top 20 annotated terms from gene ontology cellular components

### Microsatellite/Simple Sequence Repeats (SSR) prediction in Citrus reticulata

Among the variable number tandem repeats (VNTRs), microsatellites or short-sequence repeats (SSR) consist of 2-6 tandem repeats and are found in almost all eukaryotic taxa including plants, animals and humans. These markers are very useful due to their abundance, reproducibility, multiallelic nature, high genome coverage, co-dominancy and variability across the whole genome, hence these can be used for cultivar identification and parentage analysis in plants (Young and Vivier. 2010). The sequences flanking the repeats are often conserved and can be used for the primer design and the subsequent amplification of a particular SSRs.

The *in-silico* identification of microsatellite in Kinnow was performed with GMATA, which is the multi-functional tool and specifically developed for the microsatellite (SSR) mining, marker analysis and statistical calculations at the whole-genome level, it also highlights the unique genic features with SSR markers and SSR loci; accordingly, it can link the identified SSR markers with all genomics network. In the present work, we identified 111,032 SSRs markers characterized by a number of repeats ranging from 2 (di-repeats) to 6 (Hexa-repeats) bp (Table 2).

**Table 2:** SSR mining from assembled genome of *Citrus reticulata*

| Short Tandem Repeats | Total counts | Percentage |
| --- | --- | --- |
| Di-repeats | 78801 | 70.97 |
| Tri-repeats | 27115 | 24.42 |
| Tetra-repeats | 4001 | 3.60 |
| Penta-repeats | 777 | 0.69 |
| Hexa-repeats | 338 | 0.30 |
| **Total** | 111032 | 99.98 |

Further data mining on the motif types identified that dinucleotides SSR was the most common with 70.97% (78801), while trinucleotides (27115; 24.42%), tetranucleotides (4001; 3.60%), pentanucleotides (777; 0.69%), and, hexanucleotides (338; 0.30%) were progressively less common. Twelve types of di-repeats were observed with total counts of 78,801. The di-tandem repeat frequency distribution was in the following order. AT/TA (17.59%+17.45%), AG/GA (6.41%+4.23%), TC/CT (5.07%+4.48%), TG/GT (4.61%+2.83%), AC/CA (4.60%+3.08%), GC/CG (0.32%+0.28%) (Fig 4a). Similarly, sixty different types of tri-repeats were observed with total counts of 27115. The most represented tri-tandem repeats were: AAT (3.69%), ATT (3.02%), TTA (2.8%), TAA (2.29%), TAT (1.69%), ATA (1.55%), AAG (0.69), TTC (0.65%), TCT (0.51%), GAA (0.49%) (Fig 4b).

**Figure 4:**
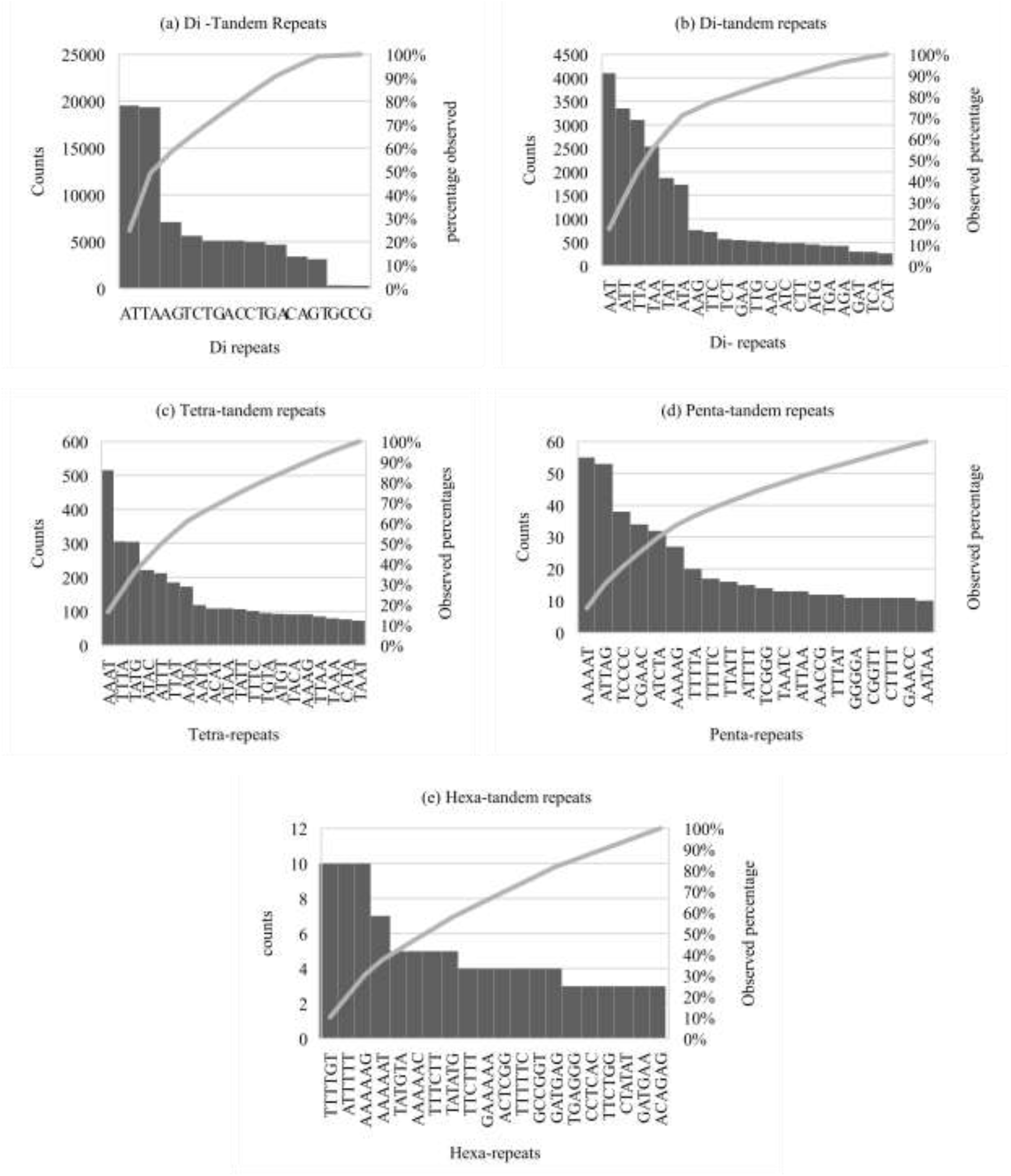
SSR distribution across the whole genome (a) Di-repeats (b) Tri-repeats (c) Tetra-repeats (d) Penta-repeats (e) Hexa-repeats

Moreover, one hundred and fifty-nine types of tetra-repeats with total counts of 4001 observed. The top five tetra-tandem repeat motif’s frequency distribution was in the following order AAAT (0.46%), TTTA (0.28%), TATG (0.27%), ATAC (0.2%), ATTT (0.19%). Fig 4 c highlight the details of observed tetra repeats with total counts and percentage. In contrast, all penta-repeats and hexa-repeats were observed at less than 1%. Details of both repeats is illustrated in figure 4-d & e respectively. One hundred and seventy-seven types of penta-repeats while 210 types of hexa-repeats were observed indicating high abundancy of hexa-repeats in present genome. Detailed of all SSRs from di to hexa-repeats are given in Supplementary Table (S8-S13).

The top contigs showing maximum no. of SSR loci are shown in Fig 5. SSR loci length ranged from 10 to 75 bp in whole genome.

**Figure 5:**
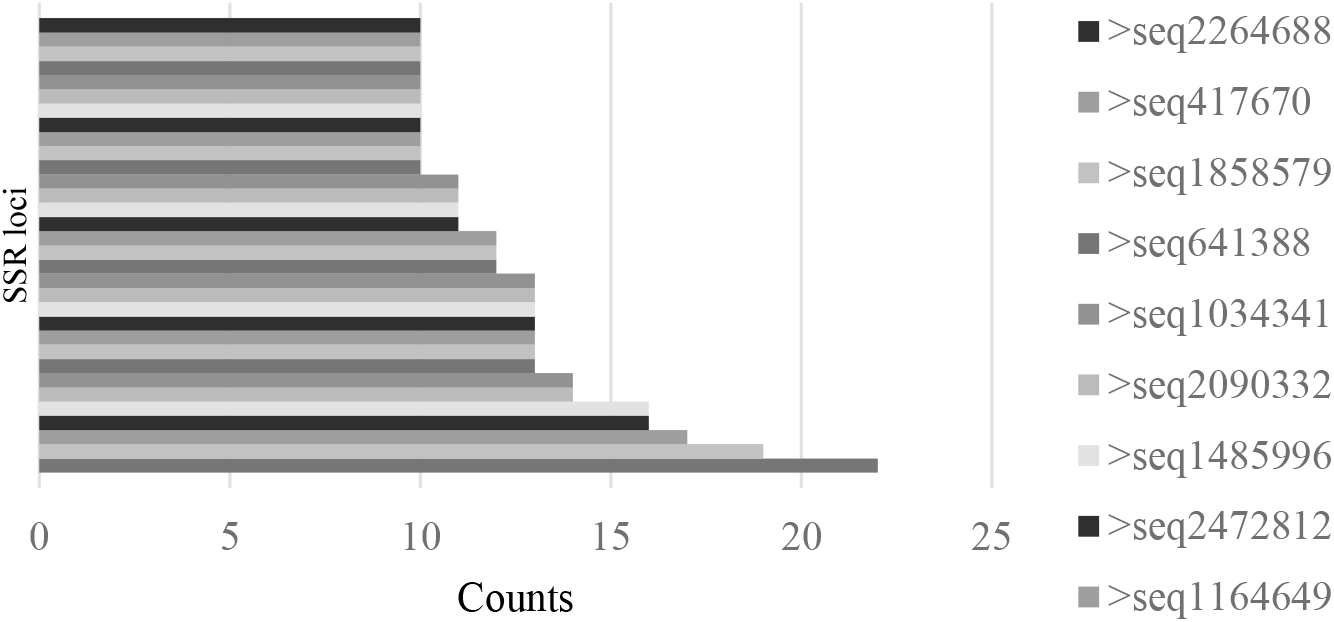
Top contigs with maximum number of loci.

For 46,531 SSRs (41.90%) the design of the primer was possible (Supplementary Table S14). The designed primers ranged in sizes from 18 to 23 base pairs with annealing temperature of 57-62°C. The GC content of all the designed primers was between 40 to 70%. The length of the end product was between 100 to 400 bp. In future, the identified unique SSR markers can be used to develop DNA barcodes of citrus species. After barcoding these markers can be used to identify the homozygosity and heterozygosity, parentage analysis, and cultivar identification in the different citrus species.

## Discussion

Whole genome sequencing of citrus species has been a major focus of plant genomics research in recent years. Citrus is an economically important crop, and sequencing the genomes of different citrus species can provide insights into their genetics and biology, as well as facilitate the development of improved citrus varieties.

Several citrus species have been sequenced using whole genome sequencing approaches. There are currently ten assembled genomes for citrus and its relatives including Clementine (*Citrus clementina*), Papeda (*Citrus ichangensis*), Sweet orange (*Citrus sinensis*), Mandarin orange (*Citrus reticulata*), Pummelo (*Citrus maxima*), Citron (*Citrus medica*), Lemon (*Citrus limon*), Hardy Orange (*Poncirus trifoliata*), Chinese Box Orange (*Atalantia buxifolia*) and Hong Kong kumquat (*Fortunella hindsii*) (Liu et al. 2022)

Illumina HiSeq 2500 platform was chosen to sequence the whole genome of Kinnow in the current study followed by *de-novo* genome assembly and functional annotation with Platanus and Hayai-Annotator Plants tools respectively. The results of genome assembly revealed the following results contigs:2630189, N50:1614 bp and GC content:34.1%, while total scaffolds were 348861 with N50 of 4827 bp. Whereas the previous assembly of Seeded Kinnow (*Citrus reticulata*) the whole genome sequencing and assembly was performed with Illumina paired end short reads. As a result, 64.2 Gb of raw data with 199.7× genome coverage was generated. The corrected and cleaned reads were assembled by plant assembler Platanus and gaps were closed by Gap Closer. The assembled scaffolds consist of 334,219,490 base pairs with N50 of 1,705,373, while the largest scaffold was observed to be 7,195,442. The length of the largest contig was 295,805 base pairs with at N50 of 24761 base pairs. In total 42653 transcripts harbor 28820 genes were predicted using homology, ab-initio and RNA sequences comparison. The average length of coding sequences was observed to be 1210 bp (Wang *et al*., 2018). Similarity among the previous and recent results suggesting that the genomic contents of Seedless Kinnow might be similar to those of the Seeded Kinnow.

SSR markers are extensively used in plants genotyping, association mapping, genetic diversities, linkage mapping and population structure analysis (Zalapa *et al*., 2012 ; Sugita *et al*., 2013 ; Gyawali *et al*., 2016 ; Zhao *et al*., 2017). In recent years several SSR markers have been developed in many crops (Cheng *et al*., 2016 ; Bhattarai and Mehlenbacher. 2017 ; KL *et al*., 2019 ; Biswas *et al*., 2020). Traditional methods for SSR screening in laboratories are time consuming, costly, and labor-intensive (Zane *et al*., 2002). The solution to this problem can be represented by in-silico detection, which are cost effective, efficient and robust (Sharma *et al*., 2007). Hence, GMATA tool was used in the current research for mining the SSRs. In total, 111032 SSR markers identified which consist of di to hexa-repeats. It was observed that as the number of nucleotides increases in repeats the frequency decreases but the types of motifs increase, e.g., di-repeats were observed to be more in counts (70.97%) but the hexa-repeats were only observed in the 0.38% of the cases. The results are compatible with the previous studies in citrus and other plants (Arabidopsis, rice, maize, soybean, tomato, cotton, and poplar) (Cardle *et al*., 2000 ; Temnykh. 2001 ; Mun *et al*., 2006 ; Biswas *et al*., 2014). A large number of e-SSR markers with primers set have been detected here and reported as (Supplementary Table S14). These markers can be used in genomic structural architecture and landscaping in all citrus species.

## Conclusion

The present research represents first time *de-novo* whole-genome assembly of Kinnow, its functional annotations and the detection of SSR loci scattered across the genome. These results will be a significant addition in the genome resources for the citrus researchers, farmers/breeders, and policy makers to propagate its valued traits among the other citrus plants regionally and globally. This genome resource will contribute to understand the evolution of various genotypes and phenotypes underlaying this unique citrus species from distinct geographical habitat.

## Credit Author Statement

Sadia Jabeen (SJ): conceptualization of the project, sample collection, *de-novo* genome assembly analysis, initial draft write-up. Rashid Saif (RS): *de-novo* genome assembly & annotations, SSR marker identification, data analysis, interpretation of results and initial draft write-up. Waseem Haider (WH): *de-novo* genome assembly design, proof-reading, Shagufta Naz (SN): conceptualization of the project, reviewing and overall supervision. Rukhama Haq (RH) conceptualization of the project, co-supervision, reviewing, editing and proofreading of the draft. Akber Hayat (AH): identification of plants, proof reading of the manuscript.

## Availability of Data

NGS-binary alignment maps (bam) files are submitted at NCBI SRA project ID: PRJNA821664. Furthermore, fourteen supplementary tables and five supplementary figures are also provided along with the manuscript.

