## Supplementary Figure 3 for "*De novo* Genome Assembly, Functional Annotation and SSR Mining of *Citrus reticulata* “Kinnow” from Pakistan"

**Shagufta Naz^1*^**

^1^Department of Biotechnology, Lahore College for Women University, Lahore, Pakistan

^2^Decode Genomics, 323-D, Punjab University Employees Housing Scheme, Lahore, Pakistan

^3^Department of Agriculture, Food and Environment, University of Catania, Italy

^4^Department of Bioscience, COMSATS University Islamabad

^5^Citrus Research Institute, Sargodha, Pakistan

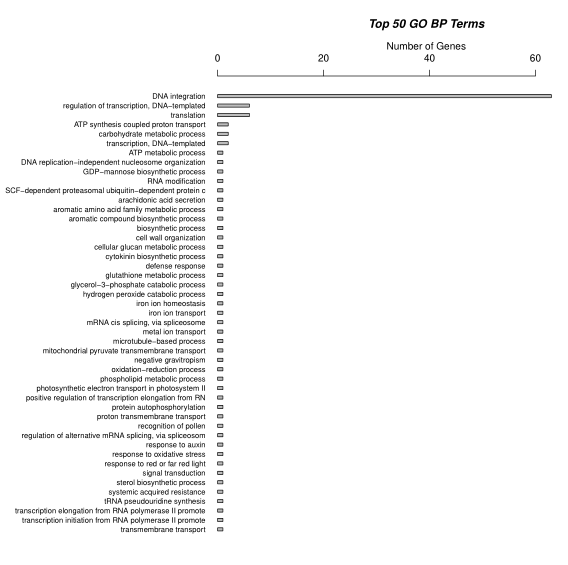

**Fig S3**: Top 50 GO:BP terms
